# Genomic selection accuracy and bias using imputed genotypes on growth, welfare and fitness traits in two Pekin duck lines

**DOI:** 10.64898/2025.12.24.696349

**Authors:** Oswald Matika, Eirini Tarsani, Kiah McIntosh, Suzanne Desire, Fasil G. Kebede, Andrea Talenti, Anne M. Rae, Andreas Kranis, Kellie A. Watson

**Affiliations:** The Roslin Institute and Royal (Dick) School of Veterinary Studies, The University of Edinburgh, Easter Bush, EH25 9RG, Midlothian, UK; Scotland’s Rural College (SRUC), Department of Animal and Veterinary Sciences, Easter Bush, EH25 9RG, Midlothian, UK; Cherry Valley Farms (UK) Ltd., Cherry Valley House, Blossom Avenue, Humberston, Grimsby. DM36 4TQ. UK; Centre for Tropical Livestock Genetics and Health (CTLGH), Roslin Institute, University of Edinburgh, Easter Bush Campus, EH25 9RG, UK

**Keywords:** Pekin ducks, imputation, genomic selection, selection accuracy, selection bias

## Abstract

The current study investigated the genomic selection accuracies and biases estimates from two commercial Pekin duck lines reared under commercial breeding practices. A large dataset of 26K duck records comprising both phenotype and imputed genotype information (60K chip) were analysed for growth, welfare and primary feather length traits. First, we employed mixed linear models with relationship matrices computed from the pedigree (BLUP) or markers (GBLUP) to estimate the variance components and breeding values. Then, we estimated the selection accuracies and selection biases to assess the more appropriate models. Our results showed moderately high imputation accuracies of 0.93 and 0.92 for lines A and D respectively. In both lines, the heritability estimates obtained using the pedigree were generally higher than using genomic markers in all traits considered. These ranged for juvenile weight (JW) from 0.22±0.01 vs 0.25±0.01 in line A vs line D using marker information to 0.39±0.02 to 0.50±0.02 using the pedigree in line A vs line D for slaughter body weight (BW). We observed very low estimates of heritability for gait 0.07±0.01 using markers in both lines. Breast muscle depth (BD) also had lower estimates of 0.15-0.16 using markers. For line A, the genomic predictions were generally higher when using the G-matrix than the A-matrix with the highest prediction was for BW (r^2^=0.68-0.70) and JW with r^2^ of 0.49. The estimates for gait and foot pad dermatitis (FPD) were greatly improved by using the G-Matrix at 0.58 vs 0.24 and 0.68 vs 0.44 respectively for markers vs pedigree information. For line D, the same improvements for G-Matrix vs A-Matrix were observed with estimates for BD being similar in the two lines. However, for BD the G-Matrix greatly improved the estimates from 0.50 to 0.71 unlike in line A where they remained at 0.50. The bias in line A were minimal (0.01- 0.19) using the G-Matrix compared to 0.02- 0.41 when using A-Matrix. The highest observed bias was for JW followed by BD for the G-matrix whereas when using the A-matrix we observed higher biases in many traits (JW, BW, BD and gait). The biases for line D were generally lower for the G-matrix (0.02 - 0.17 vs 0.00 - 0.19) than those observed in line A using markers whereas higher biases were observed using the pedigree (0.01 - 0.37). Current findings pinpointed that all traits were heritable with higher prediction accuracies and lower biases when using GBLUP as opposed to traditional BLUP.

The present study demonstrates the effectiveness of GBLUP for improving prediction accuracy and reducing bias in selection traits of Pekin ducks, particularly for traits with low heritability.

**Author Summary:** The study explored genomic selection in two commercial Pekin duck lines. Using a large dataset of 26,000 records, including phenotype and genotype data, researchers analyzed growth, welfare, and feather length traits. They applied statistical models to assess variance components and breeding values, comparing traditional pedigree-based methods (BLUP) with genomic marker-based methods (GBLUP). Results showed high imputation accuracies (93% for line A and 92% for line D). Heritability estimates varied, with genomic markers generally producing lower estimates than pedigrees, except for traits like gait and breast muscle depth where genomic predictions were superior. For example, line A showed higher accuracy using genomic data for body weight and juvenile weight. Overall, genomic predictions (GBLUP) provided higher accuracy and lower bias compared to traditional methods, especially for traits with low heritability. This highlights the effectiveness of GBLUP in improving selection processes in Pekin ducks.

## Introduction

The Pekin duck has been well documented for growth performance and meat quality traits [1–3]. In the past, poultry breeding programmes have relied on pedigree information usually gathered using single sires (ensuring the male parent is known) housed with a group of females, where artificial trap nest are used to identify the female parent. However, these are labour intensive, costly to manage and do not reflect normal poultry management practices. In addition, these may not be amenable to ever changing consumer and regulatory frame works. The use of open pens for mating allowing many males and females in a single pen has been made possible by the use of parentage panels through Single Nucleotide polymorphism (SNP) genotyping. This can be achieved through use of custom designed low density (LD) SNP genotype panels (300 to 1000 SNPs) or Medium-density (MD) SNP arrays (60k) and/or imputation. Parentage information is then used to construct pedigree information. Alternatively the marker information can directly be used to construct relationship matrices to be used in the computation of breeding values, genomic selection (GS) and overall duck improvement programmes. GS was first proposed by Meuwissen, Hayes (4), using genotypes from a reference population to predict phenotypes when only marker information is available. Since then GS has been extended to combine both genotyped and ungenotyped animals when the pedigree is available in what is termed single-step GBLUB [5]. Improved prediction accuracies have been reported in many species [6–8] when historic pedigreed data is available.

The use of G-matrices e.g. pair-wise identities by state (IBS) such as proposed by VanRaden (9) has several advantages over pedigree based A-matrix such as better estimating the Mendelian sampling term in close relatives [10–12], improving accuracies and efficiency [4, 12–14], offering the possibility of reducing pedigree-based inbreeding by using own rather than family information [11] and reducing generation interval by using younger animals for breeding [15]. In addition, genomic selection has lower selection biases estimates [16, 17] than pedigree-based selection practices.

Since genotyping using the MD SNP Chip is expensive for large flocks/herds, the strategy adopted for the current nucleus breeding flock was to use imputation from the parentage LD chip to MD Chip first proposed by [18] and used in layer chicken [19]. However, the accuracy of imputation is affected by many factors including the number of SNPs in the LD chip and their distribution along the genome, the size and connectedness of the reference population and the population genotyped with the LD Chip, marker allele frequencies etc. [20–22].

Many studies in different species e.g. sheep [13, 23–25], cattle [26, 27], aquaculture [28, 29], pigs [30, 31] and chicken [19, 32] among other species have been published on the merits of genomic selection in livestock species, the few studies published for ducks which used real data had smaller sample sizes [14, 33–35].

The traits considered in this current study include growth, primary feather length and low heritability traits for welfare such as gait as a proxy for leg health. Primary feather length (prf) has been studied in poultry species for feather growth has a significant meaning in understanding adaptive evolution, physiology, and mating of avian species. They are also important for thermoregulation. GWAS results on primary feather length in ducks identified QTL with potential for effects on growth and maturity in ducks [36, 37]. The primary feathers of ducks have been reported to have economic value of their own when transformed into products such as badminton shuttles, feather pens and more recently into complex industrial products [38–40]. On farms, they can be used as an indication of maturity.

Genetic selection for increased body weights in chicken resulted in associated problems of locomotion (gait) [41], therefore it is imperative to measure gait score in any poultry breeding programme to mitigate any potential future welfare issues [42–44].

Foot-pad dermatitis (FPD), an ulceration of the skin of the foot in poultry [45], is associated with welfare issues [46–48] and in poultry has a high reported prevalence in European countries [49]. Scoring of footpads is used as an indicator to assess welfare in broiler production systems [50] and in ducks [51].

The aim of the current study is to investigate the genomic selection accuracies and biases estimates from two commercial Pekin duck lines reared under commercial breeding practices. The parentage genotype panel (Low density panel) captured as part of a genomic parentage programme across two lines of pedigree ducks was imputed to a medium density 60K SNP panel (MD). We then estimated variance component parameters, genomic prediction accuracies and biases on 13K records per line on growth, gait and primary feather length traits. Initially, data were analysed using mixed linear models with relationship matrices computed from pedigree (BLUP) or markers (GBLUP) to estimate the variance components and breeding values. After which, we assessed the predictive genomic performance using forward prediction by estimating the selection accuracies and selection biases.

## Results

### Variance component analyses

There were high imputation accuracies of 0.93 and 0.92 for lines A and D respectively. We observed lower accuracies at the telomere and on the sex chromosome, Z.

All traits presented low to moderate estimates of heritability in either pedigree or marker information in both lines (Table 3). The estimates of heritability using the pedigree were generally higher than using genomic information in all traits considered in both lines. These ranged from 0.22±0.01 for juvenile weight (JW) vs 0.25±0.01 in line A vs line D using marker information to 0.39±0.02 to 0.50±0.02 using pedigree in line A vs line D for slaughter body weight (BW) (Table 3). Very low estimates of heritability were recorded for gait 0.07±0.01 using markers in both lines. BD had also lower estimates of 0.15-0.16 using markers

**Table 1:** Number of records used in growth trait genomic prediction for duck lines A and D.

| **Traits** | **A** | **D** |
| --- | --- | --- |
| JW | 13610 | 12964 |
| BW | 13640 | 13020 |
| BD | 11875 | 11241 |
| PRF | 11900 | 11277 |
| GAIT | 13635 | 13012 |
| ADG | 13638 | 13020 |
| FPD | 12126 | 10693 |

**Table 2.** The number of records in different generations (GEN) used in the genomic predictions for Line A and D.

| GEN | A | D |
| --- | --- | --- |
| 1 | 136 | 111 |
| 2 | 105 | 312 |
| 3 | 264 | 2780 |
| 4 | 4125 | 4932 |
| 5 | 4949 | 4885 |
| 6 | 4066 | NA |
| Total | 13645 | 13020 |

**Table 3:** Heritability estimates (h^2^) for Duck lines A and D growth using pedigree (A-Matrix) and relationship matrices (G-Matrix) using all available records.

|  | Line A |  |  | Line D |  |  |
| --- | --- | --- | --- | --- | --- | --- |
| | G-Matrix | A-Matrix | $h^2_G/h^2_A$ | G-Matrix | A-Matrix | $h^2_G/h^{2A}$ |
| JW | 0.23±0.01 | 0.41±0.02 | 0.56 | 0.25±0.01 | 0.46±0.02 | 0.54 |
| BW | 0.33±0.01 | 0.31±0.02 | 1.06 | 0.37±0.02 | 0.50±0.02 | 0.74 |
| BD | 0.17±0.01 | 0.22±0.02 | 0.77 | 0.15±0.01 | 0.21±0.02 | 0.71 |
| PRF | 0.25±0.01 | 0.31±0.02 | 0.81 | 0.19±0.01 | 0.26±0.02 | 0.73 |
| GAIT | 0.07±0.01 | 0.10±0.01 | 0.70 | 0.07±0.01 | 0.10±0.01 | 0.70 |
| ADG | 0.17±0.01 | 0.13±0.01 | 1.31 | 0.22±0.01 | 0.31±0.02 | 0.71 |
| FPD | 0.25±0.01 | 0.30±0.02 | 0.83 | 0.21±0.01 | 0.25±0.02 | 0.84 |

### Genomic predictions

The genomic predictions were generally higher using the G-matrix than the A-matrix for line A (Table 4). The highest prediction was for body weights (r^2^=0.68-0.70) outside those of juvenile weight with r^2^ of 0.49. The estimates for gait and FPD were greatly improved by using the G-Matrix 0.58 vs 0.24 and 0.68 vs 0.44 respectively for markers vs pedigree information.

**Table 4:** Genomic selection parameter estimates from Line A growth traits data obtained from generation 1 to 5 to predict generation 6 and selection bias using both A and G matrices in Asreml software with heritability estimates (h^2^) from 1-5 GEN records.

| Trait | G-Matrix |  |  |  |  | A-Matrix |  |  |  |  |
| --- | --- | --- | --- | --- | --- | --- | --- | --- | --- | --- |
| | $h^2$ | se | $r^2$ | acc | bias | $h^2$ | se | $r^2$ | acc | bias |
| JW | 0.25 | 0.02 | 0.24 | **0.49** | 0.19 | 0.44 | 0.03 | 0.16 | **0.24** | 0.41 |
| BW | 0.32 | 0.02 | 0.39 | **0.68** | -0.02 | 0.36 | 0.03 | 0.21 | **0.35** | 0.08 |
| BD | 0.20 | 0.02 | 0.22 | **0.50** | 0.11 | 0.23 | 0.02 | 0.13 | **0.26** | 0.28 |
| PRF | 0.24 | 0.02 | 0.34 | **0.69** | 0.01 | 0.31 | 0.03 | 0.22 | **0.40** | 0.06 |
| GAIT | 0.06 | 0.01 | 0.14 | **0.58** | -0.02 | 0.08 | 0.01 | 0.07 | **0.24** | 0.25 |
| ADG | 0.15 | 0.01 | 0.25 | **0.65** | -0.09 | 0.11 | 0.02 | 0.13 | **0.40** | -0.18 |
| FPD | 0.26 | 0.02 | 0.34 | **0.68** | 0.01 | 0.29 | 0.03 | 0.23 | **0.44** | 0.02 |

Where Accuracy (acc)= $\text{corr}(\text{EBV}_{\text{nopheno}}, \text{TBV}_{\text{withpheno}}) / \sqrt{h^2}$ , $b = \text{reg.corf}(\text{EBV}_{\text{nopheno}}, \text{TBV}_{\text{withpheno}})$ standard bias={1-b for $b < 1$ , $(1/b)-1$ for $b > 1$ }

The same improvements for the G-Matrix vs A-Matrix were observed in Line D (Table 5). The estimates for BD were similar in the two lines, however, the use of slaughter body weight greatly improved the G-Matrix estimates from 0.50 to 0.71 unlike in line A where they remained at 0.50.

**Table 5:**
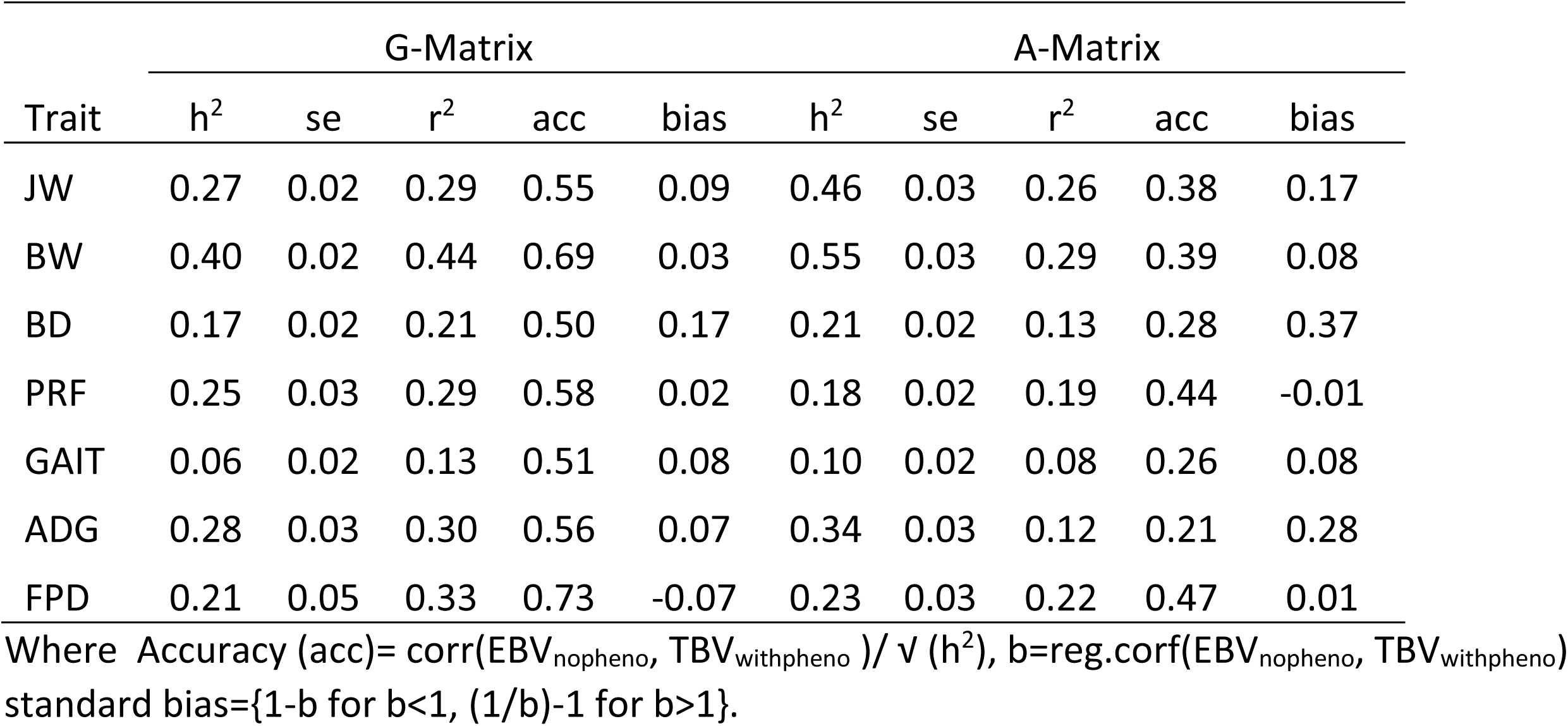
Genomic selection parameter estimates from Line D growth traits data obtained from generation 1 to 4 to predict generation 5 and selection bias using both A and G matrices in Asreml software with heritability estimates (h^2^) from 1-4 GEN records.

| Trait | G-Matrix |  |  |  |  | A-Matrix |  |  |  |  |
| --- | --- | --- | --- | --- | --- | --- | --- | --- | --- | --- |
| | $h^2$ | se | $r^2$ | acc | bias | $h^2$ | se | $r^2$ | acc | bias |
| JW | 0.27 | 0.02 | 0.29 | 0.55 | 0.09 | 0.46 | 0.03 | 0.26 | 0.38 | 0.17 |
| BW | 0.40 | 0.02 | 0.44 | 0.69 | 0.03 | 0.55 | 0.03 | 0.29 | 0.39 | 0.08 |
| BD | 0.17 | 0.02 | 0.21 | 0.50 | 0.17 | 0.21 | 0.02 | 0.13 | 0.28 | 0.37 |
| PRF | 0.25 | 0.03 | 0.29 | 0.58 | 0.02 | 0.18 | 0.02 | 0.19 | 0.44 | -0.01 |
| GAIT | 0.06 | 0.02 | 0.13 | 0.51 | 0.08 | 0.10 | 0.02 | 0.08 | 0.26 | 0.08 |
| ADG | 0.28 | 0.03 | 0.30 | 0.56 | 0.07 | 0.34 | 0.03 | 0.12 | 0.21 | 0.28 |
| FPD | 0.21 | 0.05 | 0.33 | 0.73 | -0.07 | 0.23 | 0.03 | 0.22 | 0.47 | 0.01 |

Where Accuracy (acc)= $\text{corr}(\text{EBV}_{\text{nopheno}}, \text{TBV}_{\text{withpheno}}) / \sqrt{h^2}$ , $b = \text{reg.corf}(\text{EBV}_{\text{nopheno}}, \text{TBV}_{\text{withpheno}})$ standard bias={1-b for $b < 1$ , $(1/b) - 1$ for $b > 1$ }.

The biases in line A were minimal (0.01- 0.19) when using the G-Matrix compared to 0.02-0.41 those when using the A-Matrix. The highest observed bias was for JW followed by BD for the G-matrix whereas using the A-matrix higher biases were observed in many traits (JW, BW, BD, gait etc), see Table 4 for more details. The biases for line D were generally lower for G-matrix (0.02 - 0.17 vs 0.00 - 0.19) than those observed in line A when using markers (Table 5). However, higher biases were observed using the pedigree (0.01 - 0.37).

## Discussion

The main objective of the present study was to explore forward predictive power of GS using a large dataset on accuracies and biases of the estimated breeding values on selection candidates (without own records) on both growth, welfare and fitness traits in Pekin ducks. In genomic selection, variance components of linear mixed models are ideally estimated with all available data to avoid bias. However, due to computational limitations in handling large datasets or complex models, we first fitted fixed effects and then used the residuals as phenotypes. This approach addresses both hardware and software constraints effectively. Notably, the heritability estimates obtained were within expected ranges, validating this method. Additionally, in scenarios such as across-country studies [52, 53] or when combining data from private companies, deregressed EBVs or pre-corrected phenotypes are typically used since sharing raw data can be sensitive.

Our current study is unique in that it used two different lines, over 13k records per line and seven diverse traits using imputed data to study variance components and genomic selection accuracies and biases in a commercial setting using real field data. All traits were heritable, demonstrating genetic influence across growth, welfare and fitness traits. Multidimensional plots revealed no discernible population structure, indicating a uniform genetic background within the lines (See Fig 1 and 2).

**Figure 1.**
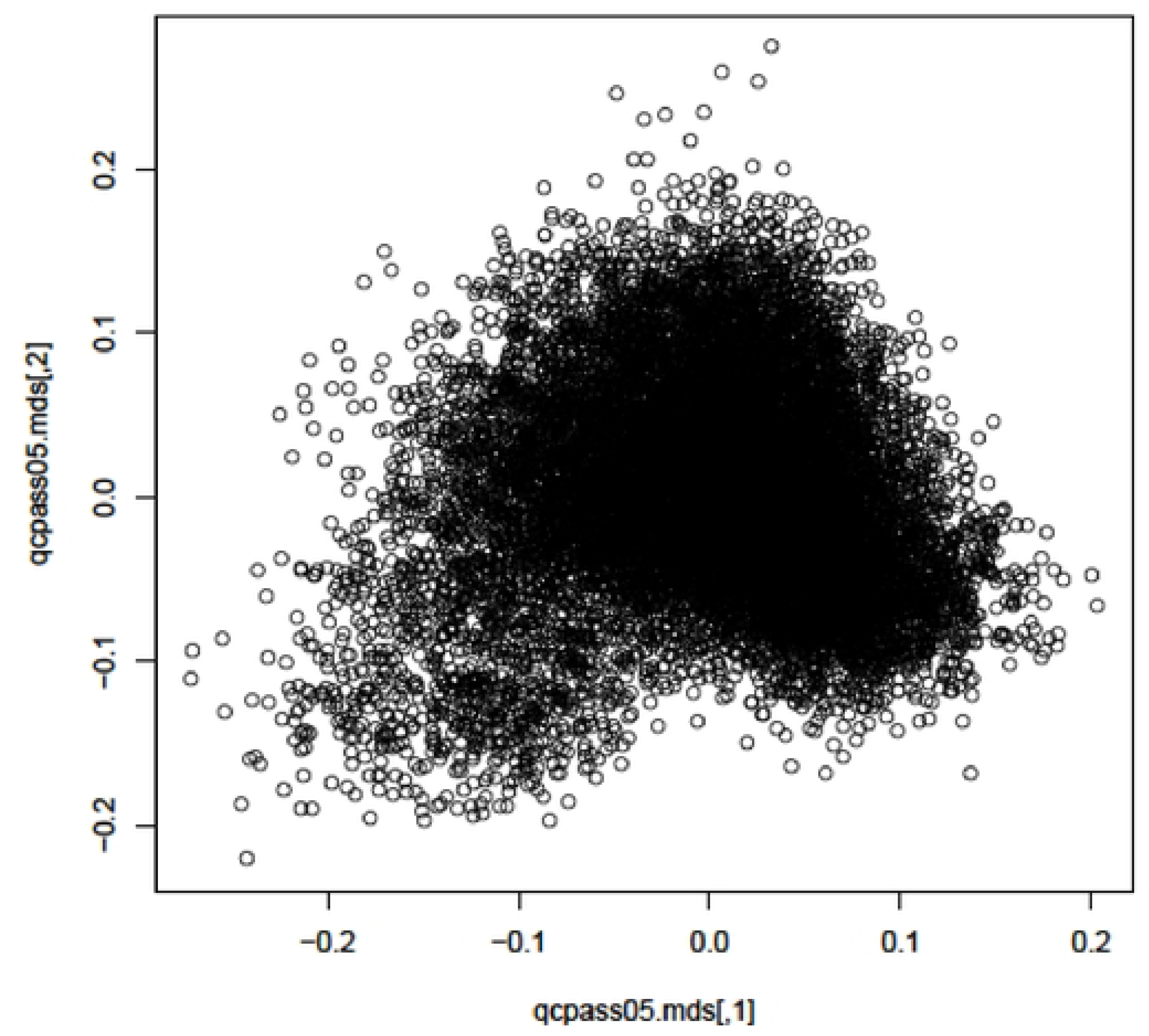
1 Multi-dimensional plot generated from 47156 Imputed SNPs on 13645 records for Line A using GenABEL software.

**Figure 2.**
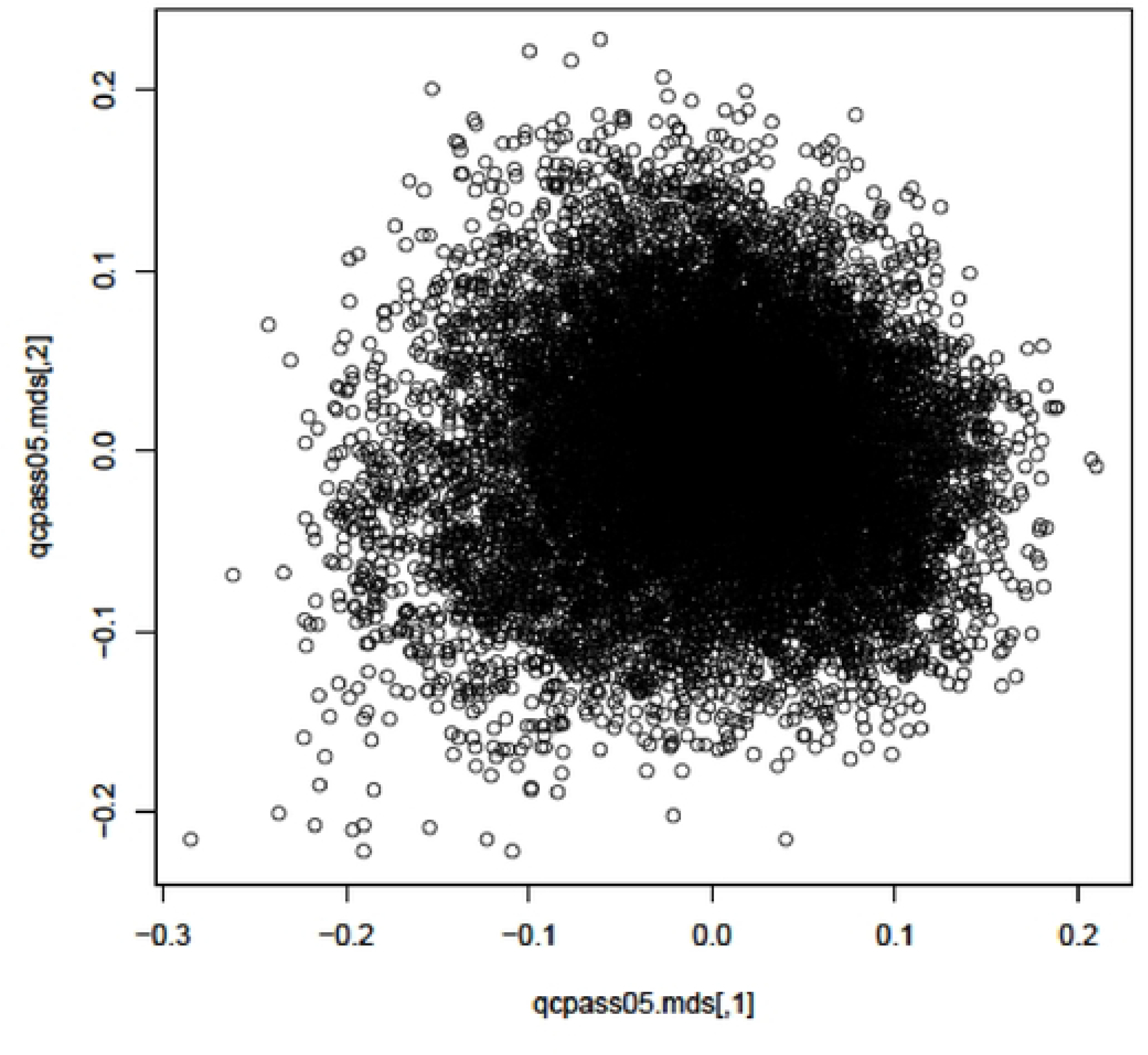
1 Multi-dimensional plot generated from 46297 imputed SNPs on 13020 records for Line D using GenABEL software.

In our study, we explored a range of traits with varying heritability estimates, from low for gait to moderate for primary feather patterns and foot pads, and generally high estimates for production traits when using the A-matrix. We observed in general, lower heritability estimates when we used the G-Matrix as opposed to the A-Matrix. The estimates of heritability were very different between the two lines. The proportion of heritability estimates recovered by the markers was also different in the two lines as we observed an overestimation in Line A, of heritability estimates in BW and ADG. However, this was not the same case for the same traits in line D, although GWAS identified a large QTL [54]. The reasons for missing heritability are many and varied and well-studied in human height and diseases [55, 56] with solutions including among others increasing sample sizes, using more rare variants and structural variations [57] not captured in SNP-chips used to create the G-Matrix [58]. Some attribute the missing heritability to rare variants that can be captured by whole sequence data [59] and use of haplotypes [60]. Génin (58) in a review, believed that the “missing heritability problem” is an ill-posed problem and may be due to complexity of the biology of traits rather than statistical methods of partition heritability estimates. Golan, Lander (61) attributed the problem of missing heritability to the use of common variants and that restricted maximum likelihood (REML) estimation underestimated the fraction of heritability due to common variation considerably. Instead, they propose phenotype correlation-genotype correlation (PCGC) regression. In their study of six diseases, they estimated the proportion of heritability due to common variants from 41% to 68% (mean 60%) using their method. Others attribute the lack of concordance between markers and traditional methods to lack of capturing additive epigenetic variances [55, 62].

Hu, Poivey (63) reported heritability of primary feather length of 0.37±0.04 in males and 0.14±0.02 in females. They also reported high genetic correlations of PRF with body weights at 10 and 18 weeks of age implying that selecting for PRF will have indirect effects on growth.

The current estimates of heritability for gait were low (0.06±0.06 and 0.12±0.06) and similar to those reported by Duggan, Rae (64) in Pekin flocks albeit in different years, different lines and with different numbers of records making this trait a good candidate for genomic selection.

The reported genetic component to FPD in different lines and breeds [49, 65, 66] ranged from low [66] to medium [49]. Kapell, Hill (65) reported in broilers heritability estimates of 0.18±0.02 to 0.24±0.02 using the pedigree and concluded that selection against FPD in a highly biosecure environment could improve the genetic merit for birds reared under commercial conditions since there was a high genetic correlation (0.78–0.82) between FPD in the pedigree and in the sib-tested environments. Ask (66) report heritability estimates for FPD in two broiler flocks of 0.21±0.03 and a genetic correlation with BW of -0.51±12.

We obtained higher prediction accuracy estimates when using the G-Matrix as opposed to the A-matrix and on the other hand, lower biases across the two lines irrespective of the large QTLs identified in Line D. The most significant enhancement in selection accuracy estimates was observed with gait score, used as a proxy for leg health. This aligns with expectations, as genomic selection is particularly effective for traits that are difficult to measure. Utilizing markers leads to higher predictive accuracies, which consequently results in greater genetic gains as outlined in the breeders’ equation. The response to selection R, can be written as R=i⋅r⋅σA

Where i is the selection intensity, r is the accuracy of prediction, and σ_A_ is the additive genetic standard deviation. The accuracy of GEBVs directly affects the reliability of selecting individuals with higher genetic potential. Increased accuracy leads to more effective selection, thereby increasing R, the expected genetic gain.

For GS to work, one constantly needs animals genotyped with an MD chip so that the link between generations is maintained. This requirement is less in this flock since all juveniles are genotyped for parentage ascertainment. In many previous studies cross-validation was used to calculate accuracies in sheep [13], in pigs [22] and in ducks [67, 68] among many others in poultry and dairy and beef cattle. The common thread is that usually, the animals are below a thousand and the genotyped animals per generation are often not sufficient to predict across generations. However, in the current study we had around 8k animals (reference set) to predict over 4k in the last generation (selection candidates). Other methodologies such as LR, have been proposed to evaluate the cross-validation tools using “partial” and “whole” data [69], however, we don’t feel this method is appropriate in the current study, since we have fewer records in earlier generations and the aim was to inform breeding practices in a commercial breeding nucleus flock where they are interested in forward predictive power of GS.

In some studies, they found that using the single step methods overestimated variance components [70] . In sheep, Macedo, Christensen (6) found that including additional information of meta founders reduced biases for milk yield but not in some traits but also observed over-dispersion when using single-step GBLUP. They also concluded that the use of additional genomic information increased the accuracies of GEBV for young rams in their population. However, in a review, Misztal, Lourenco (5) discussed conditions which need to be satisfied to make ssGBLUP unbiased and noted that is now the method of choice in large commercial breeding programmes (pigs, chicken etc) other than dairy cattle. Muir (71) showed that, in comparison of traditional BLUP and GBLUP, with both genotypes and phenotypes collected in many generations, the better was the accuracy and persistency of accuracy based on GEBV. They also showed that GEBV excelled for traits of low heritability regardless of initial equilibrium conditions and low persistence of genomic prediction under selection. However, in the current flock, all ducks are genotyped to determine parentage, making ssGBLUP unnecessary. In the current breeding programme, some of the conditions of persistence of linkage of markers across future generations will be mitigated with the addition of new animals genotyped with MD SNP chip. The traits with low heritability estimates such as gait will have the greatest benefit from use of GBLUP and genomic prediction.

## Conclusion

Our results are from a unique large dataset of almost 26k ducks with about 13k records per line in a species where there is limited published data on both growth and welfare traits. We also demonstrated reasonable accuracies using imputed genotype data. For all traits we analysed, we observed generally higher heritability estimates from the pedigree than genomic data, with higher prediction accuracies and lower biases when using GBLUP as opposed to traditional BLUP. It is important to note we observed higher accuracies in welfare traits which tend to have lower heritability estimates, hence the chance to make faster genetic progress in these hard to measure traits.

## Materials and Methods

### Ethics approval

No animal experimentation was carried out for this study. We used data (phenotypic and genomic) collected for routine animal husbandry and breeding programmes in our analyses. Data were produced and made available to us by Cherry Valley Farms (UK) Ltd company. Cherry Valley Farms (UK) Ltd is registered with the Animal & Plant Health Agency (APHA) and has an official veterinarian visit the farms once a month. Additionally, the company is part of the Poultry Health Scheme and the registration number for the farm from which the data were collected is 24/726/014.

### Animals

#### Management

Birds were reared according to standard commercial conditions using a starter feed for the first 14 days followed by a grower feed up to slaughter age. The light regime was implemented as follows: ducks had 23 hours of light on the first day, decreasing by an hour a day until 18 hours light was reached. The ducks then remained on 18 hours of light until slaughter.

Data consisted of 3000 ducks sampled on two commercial lines, genotyped using the custom designed 60K SNP array. The pedigree was reconstructed for a larger breeding population using a 427 custom built parentage SNP panel. For the two lines, all ducks with parentage SNP genotype were imputed to 60K array genotype using the dense 60K chip using Alphaimpute suite [72]. For the two lines, only ducks with both imputed genotype and phenotype were retained for subsequent analyses. Data from Line A comprised of 13645 and 13020 for line D ducks with records for juvenile weight at 12 days (JW) and at slaughter: slaughter body weight (kg) (BW); ultrasonic measured breast depth (BD) obtained using an ultrasound probe placed parallel to the keel bone at the deepest part of the breast muscle with measurements taken from the keel bone to the top of the muscle; primary feather length (PRF); gait score (Gait); average daily gain (ADG) and foot pad dermatitis score (FPD).

#### Imputation

Approximately 17,000 to 18,000 individuals per line were genotyped using a low-density (LD) SNP panel with 427 SNPs. For the training set used in imputation, 1,767 individuals from line A and 1,536 individuals from line D were genotyped with a medium-density (MD) 60K SNP panel. The complete description of the imputation process for the GWAS, which utilized one of the duck lines, can be found in reference[73].

The imputation accuracies were calculated and validated using 200 ducks per line masking their genotype from the 60K SNP chip array and after imputation, the same masked animals were compared to their imputed data to compute the accuracies.

All animals were imputed on the 60K SNP chip up to 63452 independent single nucleotide polymorphism (SNP) loci. After quality control (QC), SNPs with a minor allele frequency (MAF) less than 0.05 and those which did not meet the 1.00E-06 Hardy Weinberg Equilibrium (HWE) threshold were removed, remaining with a total of 47156 and 46297 SNPs for line A and D respectively, for further downstream analyses.

### Statistical Analysis

Data were initially analysed in SAS software to investigate the fixed effects for sex, hatching batch, age of dam and feeding pen at finishing. We also explored the interaction term between sex and hatch to give a fixed effect of “sexhatch”. The distributions of trait data values were checked for normality. Variance component analysis was conducted using linear mixed models fitting the environmental effects from the SAS models and fitting animal as random effect. The animal was fitted as random effect using either the available pedigree or the genomic relationship matrix.

Phenotypic data were pre-corrected in ASReml package [74] fitting the following model:

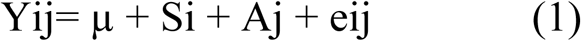

Where Yij is phenotype for JW, PRF, GAIT, FPD, Si=sexhatch and Aj=age of dam

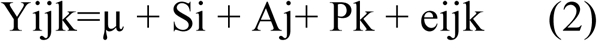

Y is phenotype for BW and BD

S=sexhatch, A=age of dam and P=feeding pen

We also fitted BW as a covariate

The resulting adjusted phenotypes or residuals (y*) were then fitted in a linear mixed model in the ASReml package [74], fitting the model:

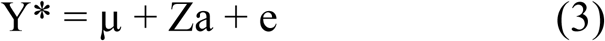

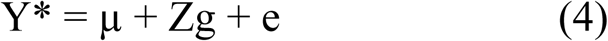

where y* is a vector of the adjusted phenotypic records, Z is a design matrix, a or g are vectors of either additive effects derived from the A-matrix or G-matrix distributed respectively as N(0, σ_a_ ^2^A) or N(0,σ_g_^2^G), σ_a_ ^2^ or σ_g_^2^ are the corresponding additive genetic variances with A and G being the relationship matrices either derived from pedigree or the markers, and e is the vector of residuals. The G matrix in the present study was constructed using the methods proposed by VanRaden (9).

Multi-dimensional plots using the markers were perform using GenABEL software [75].

### Genomic Prediction, Accuracy and Bias

Data were analysed per line (A, D) using ducks from previous generations (gen) being treated as reference (training set) and the validation sets obtained by masking the phenotype of all individuals from a given subsequent generation as will be the common practice in the commercial setting where juveniles (with no phenotype) will be selected using data from previously genotyped and phenotyped birds available. The numbers in each line are given in Table 1 and in each generation (GEN) in Table 2. For example, data for line A obtained generations 1 to 5 to predict birds in GEN 6. Genetic predictions for line D were for GEN 5 using data from GEN 1 to 4 as training set.

The genetic variance/covariance were estimated separately for each line using either markers (GEBV) or pedigree (PEBV). The reference ducks with phenotypes (non-masked) were then analysed by the model described above to predict the genomic breeding values (PGEBV) or with pedigree (PPEBV) predicted of the validation set whose phenotype were masked-phenotype individuals (i.e., In line A, 9579 ducks in generation 1 to 5, were used to predict 4066 ducks in generation 6 and in Line D, 8134 ducks from generation 1 to 4, to predict 4885 ducks in generation 5).

Pedigree or genomic prediction accuracies were calculated for each validation set (both within lines). Initially, the Pearson correlations of PGEBV with the adjusted phenotypes (rgy) were calculated and the accuracy (rgg) for each validation set was estimated by dividing rgy by the square root of the heritability of each trait for that specific validation set (in our case GEN 1-5 for line A and 1-4 for line D) as given by Legarra et al [76]:

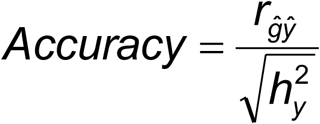

Bias was measured as the regression of “True” EBV (estimated with phenotype) on Predicted EBV (Phenotype Masked) using both marker and pedigree information. The bias (~ b) is defined as the standardised regression coefficient of TBV on EBV as explained in [16, 17] and standardised bias was given Zefreh, Doeschl-Wilson (17) in a formula below:

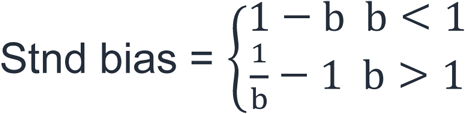

## Data Availability

The data that support the findings of this study are available from Cherry Valley Ltd upon reasonable request with signed confidentiality agreement contract by communicating with Anne Rae.

## Author Contributions

KAW, AMR and AK sourced the funding and supervised the study. OM performed all data analyses and wrote the first draft of the main manuscript. KM performed the genotype imputation and estimated the imputation accuracy. AMR collected and cured data for the study. ET, AMR, AK, SD, AT, FGK and KAW all contributed to writing and proof reading the manuscript. All authors read and approved the final manuscript.

## Corresponding author

Correspondence to Oswald Matika.

## References

1. Kokoszynski D, Wasilewski R, Steczny K, Bernacki Z, Kaczmarek K, Saleh M, et al. Comparison of growth performance and meat traits in Pekin ducks from different genotypes. European Poultry Science. 2015;79. doi: 10.1399/eps.2015.110. PubMed PMID: WOS:000369906900001.

2. Xu CC, Sun DY, Liu Y, Pan ZY, Dai ZC, Chen F, et al. Effects of ambient temperature on growth performance, slaughter traits, meat quality and serum antioxidant function in Pekin duck. Frontiers in Veterinary Science. 2024;11. doi: 10.3389/fvets.2024.1363355. PubMed PMID: WOS:001187595600001.

3. Kokoszynski D, Piwczynski D, Arpásová H, Hrncar C, Saleh M, Wasilewski R. A comparative study of carcass characteristics and meat quality in genetic resources Pekin ducks and commercial crossbreds. Asian-Australasian Journal of Animal Sciences. 2019;32(11):1753–62. doi: 10.5713/ajas.18.0790. PubMed PMID: WOS:000492677500012.

4. Meuwissen THE, Hayes BJ, Goddard ME. Prediction of total genetic value using genome-wide dense marker maps. Genetics. 2001;157(4):1819–29. PubMed PMID: WOS:000168223400036.

5. Misztal I, Lourenco D, Legarra A. Current status of genomic evaluation. Journal of Animal Science. 2020;98(4). doi: 10.1093/jas/skaa101. PubMed PMID: WOS:000530722100025.

6. Macedo FL, Christensen OF, Astruc JM, Aguilar I, Masuda Y, Legarra A. Bias and accuracy of dairy sheep evaluations using BLUP and SSGBLUP with metafounders and unknown parent groups. Genetics Selection Evolution. 2020;52(1). doi: 10.1186/s12711-020-00567-1. PubMed PMID: WOS:000563282400002.

7. Asadollahi H, Mahyari SA, Torshizi RV, Emrani H, Ehsani A. Bias and accuracy of body weight trait evaluations of an F2 chicken using single-step genomic best linear unbiased prediction (ssGBLUP). Canadian Journal of Animal Science. 2023. doi: 10.1139/cjas-2023-00091. PubMed PMID: WOS:001088383000001.

8. Alvarenga AB, Veroneze R, Oliveira HR, Marques DBD, Lopes PS, Silva FF, et al. Comparing Alternative Single-Step GBLUP Approaches and Training Population Designs for Genomic Evaluation of Crossbred Animals. Frontiers in Genetics. 2020;11. doi: 10.3389/fgene.2020.00263. PubMed PMID: WOS:000529970600001.

9. VanRaden PM. Efficient Methods to Compute Genomic Predictions. Journal of Dairy Science. 2008;91(11):4414–23. doi: 10.3168/jds.2007-0980. PubMed PMID: WOS:000260277200035.

10. Patry C, Ducrocq V. Evidence of biases in genetic evaluations due to genomic preselection in dairy cattle. Journal of Dairy Science. 2011;94(2):1011–20. doi: 10.3168/jds.2010-3804. PubMed PMID: WOS:000286407600047.

11. Liu HM, Sorensen AC, Meuwissen THE, Berg P. Allele frequency changes due to hitch-hiking in genomic selection programs. Genetics Selection Evolution. 2014;46. doi: 10.1186/1297-9686-46-8. PubMed PMID: WOS:000333516800001.

12. Clark SA, Hickey JM, Daetwyler HD, van der Werf JHJ. The importance of information on relatives for the prediction of genomic breeding values and the implications for the makeup of reference data sets in livestock breeding schemes. Genetics Selection Evolution. 2012;44. doi: 10.1186/1297-9686-44-4. PubMed PMID: WOS:000302059000001.

13. Riggio V, Abdel-Aziz M, Matika O, Moreno CR, Carta A, Bishop SC. Accuracy of genomic prediction within and across populations for nematode resistance and body weight traits in sheep. Animal. 2014;8(4):520–8. doi: 10.1017/s1751731114000081. PubMed PMID: WOS:000342206000002.

14. Zhou J, Yu JZ, Zhu MY, Yang FX, Hao JP, He Y, et al. Optimizing Breeding Strategies for Pekin Ducks Using Genomic Selection: Genetic Parameter Evaluation and Selection Progress Analysis in Reproductive Traits. Applied Sciences-Basel. 2025;15(1). doi: 10.3390/app15010194. PubMed PMID: WOS:001393448300001.

15. Pryce JE, Daetwyler HD. Designing dairy cattle breeding schemes under genomic selection: a review of international research. Animal Production Science. 2012;52(2-3):107–14. doi: 10.1071/an11098. PubMed PMID: WOS:000301064800006.

16. Lipschutz-Powell D, Woolliams J, Bijma P, Pong-Wong R, Bermingham M, Doeschl-Wilson A. Bias, Accuracy, and Impact of Indirect Genetic Effects in Infectious Diseases. Frontiers in Genetics. 2012;3. doi: 10.3389/fgene.2012.00215.

17. Zefreh MG, Doeschl-Wilson AB, Riggio V, Matika O, Pong-Wong R. Exploring the value of genomic predictions to simultaneously improve production potential and resilience of farmed animals. Frontiers in Genetics. 2023;14. doi: 10.3389/fgene.2023.1127530. PubMed PMID: WOS:001000448800001.

18. Habier D, Fernando RL, Dekkers JCM. Genomic Selection Using Low-Density Marker Panels. Genetics. 2009;182(1):343–53. doi: 10.1534/genetics.108.100289. PubMed PMID: WOS:000270213800029.

19. Herry F, Druet DP, Hérault F, Varenne A, Burlot T, Le Roy P, et al. Interest of using imputation for genomic evaluation in layer chicken. Poultry Science. 2020;99(5):2324–36. doi: 10.1016/j.psj.2020.01.004. PubMed PMID: WOS:000532702800003.

20. Calus MPL, Bouwman AC, Hickey JM, Veerkamp RF, Mulder HA. Evaluation of measures of correctness of genotype imputation in the context of genomic prediction: a review of livestock applications. Animal. 2014;8(11):1743–53. doi: 10.1017/s1751731114001803. PubMed PMID: WOS:000344117600001.

21. Marchini J, Howie B. Genotype imputation for genome-wide association studies. Nature Reviews Genetics. 2010;11(7):499–511. doi: 10.1038/nrg2796. PubMed PMID: WOS:000278998500012.

22. Badke YM, Bates RO, Ernst CW, Fix J, Steibel JP. Accuracy of Estimation of Genomic Breeding Values in Pigs Using Low-Density Genotypes and Imputation. G3-Genes Genomes Genetics. 2014;4(4):623–31. doi: 10.1534/g3.114.010504. PubMed PMID: WOS:000334694100007.

23. Rupp R, Mucha S, Larroque H, McEwan J, Conington J. Genomic application in sheep and goat breeding. Animal Frontiers. 2016;6(1):39–44. doi: 10.2527/af.2016-0006. PubMed PMID: WOS:000457256200006.

24. Lillehammer M, Sonesson AK, Klemetsdal G, Blichfeldt T, Meuwissen THE. Genomic selection strategies to improve maternal traits in Norwegian White Sheep. Journal of Animal Breeding and Genetics. 2020;137(4):384–94. doi: 10.1111/jbg.12475. PubMed PMID: WOS:000522495800001.

25. Shumbusho F, Raoul J, Astruc JM, Palhiere I, Lemarié S, Fugeray-Scarbel A, et al. Economic evaluation of genomic selection in small ruminants: a sheep meat breeding program. Animal. 2016;10(6):1033–41. doi: 10.1017/s1751731115002049. PubMed PMID: WOS:000377126700018.

26. Bouquet A, Juga J. Integrating genomic selection into dairy cattle breeding programmes: a review. Animal. 2013;7(5):705–13. doi: 10.1017/s1751731112002248. PubMed PMID: WOS:000316818500001.

27. Cole JB, da Silva M. Genomic selection in multi-breed dairy cattle populations. Revista Brasileira De Zootecnia-Brazilian Journal of Animal Science. 2016;45(4):195–202. doi: 10.1590/s1806-92902016000400008. PubMed PMID: WOS:000377019100008.

28. Zenger KR, Khatkar MS, Jones DB, Khalilisamani N, Jerry DR, Raadsma HW. Genomic Selection in Aquaculture: Application, Limitations and Opportunities With Special Reference to Marine Shrimp and Pearl Oysters. Frontiers in Genetics. 2019;9. doi: 10.3389/fgene.2018.00693. PubMed PMID: WOS:000456573800001.

29. Yáñez JM, Barría A, López ME, Moen T, Garcia BF, Yoshida GM, et al. Genome-wide association and genomic selection in aquaculture. Reviews in Aquaculture. 2023;15(2):645–75. doi: 10.1111/raq.12750. PubMed PMID: WOS:000888943300001.

30. Lillehammer M, Meuwissen THE, Sonesson AK. Genomic selection for maternal traits in pigs. Journal of Animal Science. 2011;89(12):3908–16. doi: 10.2527/jas.2011-4044. PubMed PMID: WOS:000297368200007.

31. Lillehammer M, Meuwissen THE, Sonesson AK. Genomic selection for two traits in a maternal pig breeding scheme. Journal of Animal Science. 2013;91(7):3079–87. doi: 10.2527/jas.2012-5113. PubMed PMID: WOS:000320856200011.

32. Wolc A, Kranis A, Arango J, Settar P, Fulton JE, O’Sullivan NP, et al. Implementation of genomic selection in the poultry industry. Animal Frontiers. 2016;6(1):23–31. doi: 10.2527/af.2016-0004. PubMed PMID: WOS:000457256200004.

33. Cai WT, Hu J, Fan WL, Xu YX, Tang J, Xie M, et al. Genetic parameters and genomic prediction of growth and breast morphological traits in a crossbreed duck population. Evolutionary Applications. 2024;17(2). doi: 10.1111/eva.13638. PubMed PMID: WOS:001157622700001.

34. Chen H, Luo KY, Wang C, Xuan R, Zheng SM, Tang HB, et al. Genomic characteristics and selection signals of Zhongshan ducks. Animal. 2023;17(5). doi: 10.1016/j.animal.2023.100797. PubMed PMID: WOS:000989218300001.

35. Cai W, Hu J, Fan W, Xu Y, Tang J, Xie M, et al. Genetic parameters and genomic prediction of growth and breast morphological traits in a crossbreed duck population. Evolutionary Applications. 2024;17(2). doi: 10.1111/eva.13638. PubMed PMID: WOS:001157622700001.

36. Ma SC, Li PC, Liu HH, Xi Y, Xu Q, Qi JJ, et al. Genome-wide association analysis of the primary feather growth traits of duck: identification of potential Loci for growth regulation. Poultry Science. 2023;102(1). doi: 10.1016/j.psj.2022.102243. PubMed PMID: WOS:000893117100012.

37. Wang HZ, Twumasi G, Xu Q, Xi Y, Qi JJ, Yang Z, et al. Identification of candidate genes associated with primary feathers of tianfu nonghua ducks based on Genome-wide association studies. Poultry Science. 2024;103(9). doi: 10.1016/j.psj.2024.103985. PubMed PMID: WOS:001266390300001.

38. Kurien RA, Biju A, Raj AK, Chacko A, Joseph B, Koshy CP, et al. Comparative Mechanical Properties of Duck Feather-Jute Fiber Reinforced Hybrid Composites. Transactions of the Indian Institute of Metals. 2023;76(9):2575–80. doi: 10.1007/s12666-023-03015-y. PubMed PMID: WOS:001020211900001.

39. Alvarez S, Raydan ND, Svahn I, Gontier E, Rischka K, Charrier B, et al. Assessment and Characterization of Duck Feathers as Potential Source of Biopolymers from an Upcycling Perspective. Sustainability. 2023;15(19). doi: 10.3390/su151914201. PubMed PMID: WOS:001081969100001.

40. Wang H, Jin XY, Wu HB. Adsorption and desorption properties of modified feather and feather/polypropylene melt-blown filter cartridge of lead ion (Pb2+). Journal of Industrial Textiles. 2016;46(3):852–67. doi: 10.1177/1528083715598896. PubMed PMID: WOS:000382851800011.

41. Paxton H, Daley MA, Corr SA, Hutchinson JR. The gait dynamics of the modern broiler chicken: a cautionary tale of selective breeding. Journal of Experimental Biology. 2013;216(17):3237–48. doi: 10.1242/jeb.080309. PubMed PMID: WOS:000322955200018.

42 Duggan BM, Hocking PM, Clements DN. Gait in ducks (*Anas platyrhynchos*) and chickens (*Gallus gallus*) - similarities in adaptation to high growth rate. Biology Open. 2016;5(8):1077–85. doi: 10.1242/bio.018614. PubMed PMID: WOS:000382304400008.

43. Makagon MM, Woolley R, Karcher DM. Assessing the waddle: An evaluation of a 3-point gait score system for ducks. Poultry Science. 2015;94(8):1729–34. doi: 10.3382/ps/pev151. PubMed PMID: WOS:000358186100002.

44. Robison CI, Rice M, Makagon MM, Karcher DM. Duck gait: Relationship to hip angle, bone ash, bone density, and morphology. Poultry Science. 2015;94(5):1060–7. doi: 10.3382/ps/pev050. PubMed PMID: WOS:000353347200029.

45. Mayne RK. A review of the aetiology and possible causative factors of foot pad dermatitis in growing turkeys and broilers. Worlds Poultry Science Journal. 2005;61(2):256–67. doi: 10.1079/wps200458. PubMed PMID: WOS:000229873100006.

46. Clark S, Hansen G, McLean P, Bond P, Wakeman W, Meadows R, et al. Pododermatitis in turkeys. Avian Diseases. 2002;46(4):1038–44. doi: 10.1637/0005-2086(2002)046[1038:Pit]2.0.Co2. PubMed PMID: WOS:000179736700035.

47. Martland MF. ULCERATIVE DERMATITIS IN BROILER-CHICKENS - THE EFFECTS OF WET LITTER. Avian Pathology. 1985;14(3):353–64. doi: 10.1080/03079458508436237. PubMed PMID: WOS:A1985APA4200006.

48. Rochlitz I, Broom DM. The welfare of ducks during foie gras production. Animal Welfare. 2017;26(2):135–49. doi: 10.7120/09627286.26.2.135. PubMed PMID: WOS:000401903400001.

49. Kjaer JB, Su G, Nielsen BL, Sorensen P. Foot pad dermatitis and hock burn in broiler chickens and degree of inheritance. Poultry Science. 2006;85(8):1342–8. doi: 10.1093/ps/85.8.1342. PubMed PMID: WOS:000239339300003.

50. Heitmann S, Stracke J, Petersen H, Spindler B, Kemper N. First approach validating a scoring system for foot-pad dermatitis in broiler chickens developed for application in practice. Preventive Veterinary Medicine. 2018;154:63–70. doi: 10.1016/j.prevetmed.2018.03.013. PubMed PMID: WOS:000433015000009.

51. Klambeck L, Stracke J, Spindler B, Klotz D, Wohlsein P, Schön HG, et al. First approach to validate a scoring system to assess footpad dermatitis in Pekin ducks. European Poultry Science. 2019;83. doi: 10.1399/eps.2019.262. PubMed PMID: WOS:000460544400001.

52. Sallam M, Benhajali H, Savoia S, de Koning D, Strandberg E. Across-countries genomic prediction using national breeding values or multitrait across-countries evaluation breeding values. Journal of Dairy Science. 2022;105(4):3282–95. doi: 10.3168/jds.2021-20877. PubMed PMID: WOS:000821075700022.

53. Ostersen T, Christensen O, Henryon M, Nielsen B, Su G, Madsen P. Deregressed EBV as the response variable yield more reliable genomic predictions than traditional EBV in pure-bred pigs. Genetics Selection Evolution. 2011;43. doi: 10.1186/1297-9686-43-38. PubMed PMID: WOS:000297995800001.

54. Tarsani E, Matika O, McIntosh K, Desire S, Dunn IC, Talenti A, et al. Leveraging genome-wide association analyses with chip and imputed data emerges potential pleiotropic region for four duck growth traits. Scientific Reports. 2025;submitted.

55. Bourrat P, Lu QY. Dissolving the Missing Heritability Problem. Philosophy of Science. 2017;84(5):1055–67. doi: 10.1086/694007. PubMed PMID: WOS:000418037100022.

56. Marian AJ. Elements of ’missing heritability’. Current Opinion in Cardiology. 2012;27(3):197–201. doi: 10.1097/HCO.0b013e328352707d. PubMed PMID: WOS:000302816400001.

57. Theunissen F, Flynn LL, Anderton RS, Mastaglia F, Pytte J, Jiang L, et al. Structural Variants May Be a Source of Missing Heritability in sALS. Frontiers in Neuroscience. 2020;14. doi: 10.3389/fnins.2020.00047. PubMed PMID: WOS:000515635300001.

58. Génin E. Missing heritability of complex diseases: case solved? Human Genetics. 2020;139(1):103–13. doi: 10.1007/s00439-019-02034-4. PubMed PMID: WOS:000511692000010.

59. Young AI. Discovering missing heritability in whole-genome sequencing data. Nature Genetics. 2022;54(3):224–6. doi: 10.1038/s41588-022-01012-3. PubMed PMID: WOS:000767015300003.

60. Shirali M, Knott SA, Pong-Wong R, Navarro P, Haley CS. Haplotype Heritability Mapping Method Uncovers Missing Heritability of Complex Traits. Scientific Reports. 2018;8. doi: 10.1038/s41598-018-23307-4. PubMed PMID: WOS:000427926500026.

61. Golan D, Lander ES, Rosset S. Measuring missing heritability: Inferring the contribution of common variants. Proceedings of the National Academy of Sciences of the United States of America. 2014;111(49):E5272–E81. doi: 10.1073/pnas.1419064111. PubMed PMID: WOS:000345921500005.

62. Trerotola M, Relli V, Simeone P, Alberti S. Epigenetic inheritance and the missing heritability. Human Genomics. 2015;9. doi: 10.1186/s40246-015-0041-3. PubMed PMID: WOS:000358486100001.

63. Hu YH, Poivey JP, Rouvier R, Wang CT, Tai C. Heritabilities and genetic correlations of body weights and feather length in growing Muscovy selected in Taiwan. British Poultry Science. 1999;40(5):605–12. doi: 10.1080/00071669986972. PubMed PMID: WOS:000084834200007.

64. Duggan BM, Rae AM, Clements DN, Hocking PM. Higher heritabilities for gait components than for overall gait scores may improve mobility in ducks. Genetics Selection Evolution. 2017;49. doi: 10.1186/s12711-017-0317-2. PubMed PMID: WOS:000400426700001.

65. Kapell D, Hill WG, Neeteson AM, McAdam J, Koerhuis ANM, Avendaño S. Genetic parameters of foot-pad dermatitis and body weight in purebred broiler lines in 2 contrasting environments. Poultry Science. 2012;91(3):565–74. doi: 10.3382/ps.2011-01934. PubMed PMID: WOS:000300616800004.

66. Ask B. Genetic variation of contact dermatitis in broilers. Poultry Science. 2010;89(5):866–75. doi: 10.3382/ps.2009-00496. PubMed PMID: WOS:000276893800003.

67. Zhang F, Zhu F, Yang FX, Hao JP, Hou ZC. Genomic selection for meat quality traits in Pekin duck. Animal Genetics. 2022;53(1):94–100. doi: 10.1111/age.13157. PubMed PMID: WOS:000723126900001.

68. Cai WT, Hu J, Fan WL, Xu YX, Tang J, Xie M, et al. Strategies to improve genomic predictions for 35 duck carcass traits in an F2 population. Journal of Animal Science and Biotechnology. 2023;14(1). doi: 10.1186/s40104-023-00875-8. PubMed PMID: WOS:000983214000001.

69. Legarra A, Reverter A. Semi-parametric estimates of population accuracy and bias of predictions of breeding values and future phenotypes using the LR method. Genetics Selection Evolution. 2018;50. doi: 10.1186/s12711-018-0426-6. PubMed PMID: WOS:000449386800002.

70. Wang L, Janss LL, Madsen P, Henshall J, Huang CH, Marois D, et al. Effect of genomic selection and genotyping strategy on estimation of variance components in animal models using different relationship matrices. Genetics Selection Evolution. 2020;52(1). doi: ARTN 31 10.1186/s12711-020-00550-w. PubMed PMID: WOS:000546855200001.

71. Muir WM. Comparison of genomic and traditional BLUP-estimated breeding value accuracy and selection response under alternative trait and genomic parameters. Journal of Animal Breeding and Genetics. 2007;124(6):342–55. doi: DOI 10.1111/j.1439-0388.2007.00700.x. PubMed PMID: WOS:000251432800004.

72. Cleveland MA, Hickey JM. Practical implementation of cost-effective genomic selection in commercial pig breeding using imputation. Journal of Animal Science. 2013;91(8):3583–92. doi: 10.2527/jas.2013-6270. PubMed PMID: WOS:000322588100011.

73. Tarsani E, Matika O, McIntosh K, Desire S, Dunn I, Talenti A, et al. Leveraging genome-wide association analyses with chip and imputed data emerges potential pleiotropic region for four duck growth traits. Scientific Reports. 2025;15(1). doi: 10.1038/s41598-025-08852-z. PubMed PMID: WOS:001522021300001.

74. Gilmour AR, Gogel BJ, Cullis BR, Thompson R. ASReml User Guide Release 3.0. VSN Int Ltd. 2009.

75. Aulchenko Y, Ripke S, Isaacs A, Van Duijn C. GenABEL: an R library for genome-wide association analysis. Bioinformatics. 2007;23(10):1294–6. doi: 10.1093/bioinformatics/btm108. PubMed PMID: WOS:000247348300017.

76. Legarra A, Robert-Granie C, Manfredi E, Elsen JM. Performance of genomic selection in mice. Genetics. 2008;180(1):611–8. doi: 10.1534/genetics.108.088575. PubMed PMID: WOS:000259758500049.

